# Identifying Simultaneous Rearrangements in Cancer Genomes

**DOI:** 10.1101/164855

**Authors:** Layla Oesper, Simone Dantas, Benjamin J. Raphael

## Abstract

The traditional view of cancer evolution states that a cancer genome accumulates a *sequential* ordering of mutations over a long period of time. However, in recent years it has been suggested that a cancer genome may instead undergo a one-time catastrophic event, such as *chromothripsis*, where a large number of mutations instead occur *simultaneously*. A number of potential signatures of chromothripsis have been proposed. In this work we provide a rigorous formulation and analysis of the “ability to walk the derivative chromosome” signature originally proposed by Korbel and Campbell (2013). In particular, we show that this signature, as originally envisioned, may not always be present in a chromothripsis genome and we provide a precise quantification of under what circumstances it would be present. We also propose a variation on this signature, the *H/T alternating fraction*, which allows us to overcome some of the limitations of the original signature. We apply our measure to both simulated data and a previously analyzed real cancer dataset and find that the H/T alternating fraction may provide useful signal for distinguishing genomes having acquired mutations simultaneously from those acquired in a sequential fashion. An implementation of the H/T alternating fraction is available at https://bitbucket.org/oesperlab/ht-altfrac.

## 1 Introduction

The development of cancer is driven by the accumulation of somatic mutations within a set of cells. These mutations can vary from single nucleotide variants (SNVs) to large-scale structural variations (SVs) – including deletions, duplications, inversions and translocations – that rearrange entire segments of DNA. While the traditional view of cancer evolution states that a cancer genome accumulates a *sequential* ordering of these mutations over a long period of time [17], in 2011 Stephens *et al*. [21] proposed an alternative model where instead a collection of rearrangements occur *simultaneously*. Specifically, they characterized *chromothripsis* as an event where a portion of a chromosome (or several chromosomes) shatter and a subset of the fragments are randomly stitched together. As a result, the participating chromosomes appear highly rearranged – containing a number of closely located and overlapping SVs.

While the exact underlying mechanisms that drive chromosome shattering in cancer remain unknown, a number of studies have reported the presence of chromothripsis in numerous cancer types including bone, colorectal, and prostate cancers [5, 9, 21]. Often times samples labeled as chromothripsis are associated with a poor outcome for the patient [10, 16]. Chromothripsis has also been reported in germline samples – usually in conjunction with other complex diseases [3, 8]. Furthermore, other related phenomenons have been suggested that also indicate the accumulation of multiple rearrangements at once. For example, *chromoplexy* [1] is similar to chromothripsis but generally tends to have few aberrations that affect more chromosomes. On the other hand, [12] suggested using the term *chromoanasynthesis* instead of chromothripsis in order to better reflect potential mechanisms underlying the phenomenon, but some treat this a separate category of simultaneous events [3].

A fundamental question underlying these continuing reports of chromothripsis, and other simultaneous events, is: How does one distinguish a *sequential* accumulation of mutations from a *simultaneous* acquisition? The original chromothripsis paper [21] first noted that the genomes they believed to have undergone chromothripsis exhibited some striking patterns: (1) clustering of break points along affected portions of the genome; and (2) oscillating copy number states. The later observation is a result of genomic segments in the shattered region not being included in the derivative chromosome, and therefore appearing as deleted. The initial discovery [21] and later reports of chromothripsis [14, 19] rely on Monte Carlo simulations to argue that sequential rearrangements were unlikely to produce these patterns.

Subsequently, Korbel and Campbell [11] built upon these observations and proposed a list of six criteria for chromothripsis: clustering of breakpoints, oscillating copy number states, interspersed loss and retention of heterozygosity, breakpoints affecting one haplotype, randomness amongst fragment join types and ability to walk the derivative chromosome. While approaches designed to detect complex rearrangements in cancer such as CouGaR [4], PREGO [18] or McPherson *et al*. [15] might be useful for detecting and analyzing chromothripsis, methods designed specifically for this task, including many that rely on a subset of these proposed signatures, may be more appropriate [1, 2, 6, 22]. For instance, ShatterProof [6] combines scores across several criteria to identify regions of a genome that are likely to have undergone a chromothripsis event. Cai *et al*. [2] identify what they call “chromothripsis-like patterns” or CTLP by clustering copy number status changes (effectively merging two of the signatures). So, while some of the signatures proposed by Korbel and Campbell (in particular those also reported in the original chromothripsis publication) appear to be used frequently in practice, further analysis of the other signatures may provide additional insight.

In this paper we perform an analysis of one of the signatures proposed by Korbel and Campbell, the ability to walk the derivative chromosome. In particular, we show that this signature, as originally envisioned, may not always be present in a chromothripsis genome and we provide a precise quantification of under what circumstances it would be present. We also propose a variation on this signature, the *H/T alternating fraction*, which overcomes some of the limitations of the original signature. We apply our measure to both simulated data and a previously analyzed real cancer dataset and find that the H/T alternating fraction may be useful for distinguishing genomes having acquired mutations simultaneously from those acquired in a sequential fashion.

## 2 Methods

### 2.1 Chromothripsis Strings

Consider a unichromosomal reference genome *G* where we label consecutive intervals, or genomic segments along *G* using the characters 1, 2, *…, n*. Let *S* = {1, 2, *…, n*} be the set of these characters. An interval *g* ∈ *S* that appears in the reverse orientation in a genome derived from *G* will be denoted using its *inverse –g*. We define *S̃*= {±1, …, ±*n*} to be the set of all characters in *S* and their inverses. A linear cancer genome *C* derived from *G* via genomic rearrangements is a sequence of characters from *S̃*. Therefore, *C* ∈ *S̃*∗, the Kleene closure of *S̃* (representing the set of all *rearrangement strings* that can be derived from *G*). A chromothripsis event rearranges genomic segments, allowing for deletion of some segments, but no duplication of segments. Therefore, only a subset of *S̃*∗ can be the result of a chromothripsis event. We now define which rearrangement strings in *S̃*∗ may result from a chromothripsis event.

#### Definition 2.1

*We define a linear string C ∈ S̃∗ to be a* chromothripsis string *if C is a signed permutation of 2 or more characters from S*.

### 2.2 Extremities, Adjacencies and Extremity Permutations

Each genomic interval *g* ∈ *S* can be denoted as an interval with two extremities: [*g_t_, g_h_
*] where *g_t_
* is the *tail extremity* and *g_h_
* is the *head extremity* of the interval denoted by the character *g*. We define the *extremity set E* = *T* ∪ *H* where *T* = {*g_t_
* | *g* ∈ *S*}and *H* = {*g_h_ | g* ∈ *S*}. An interval *g* ∈ *S* that appears in the reverse orientation is denoted as –*g* = [–*g_t_, –g_h_
*] = [*g_h_, g_t_
*]. Therefore, any interval *g* ∈ *S̃* can be written as an ordered pair of extremities from *E* where one extremity is from *H* and the other from *T*. Notice that once a single extremity for an interval *g* is defined, the other extremity, or *obverse extremity* is completely predetermined. Therefore, we define an *adjacency* to be an unordered pair of extremities from *E*, indicating an adjacency between two intervals from *S̃*. Suppose *g, g’* ∈ *S̃*, the adjacency between (*g, g’*) is defined in Equation (1).

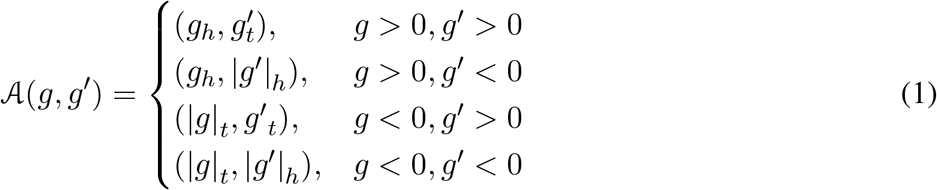

Suppose that *C* = *c*
_1_
*c*
_2_ … *c_m_
* and *C* ∈ *S̃*
^
*∗*
^. We define the *adjacency set* of *C* as *A*(*C*) = {*A* (*c_j_, c_j_
*
_+1_): *j* = 1, *…, m* 1}. We note that this manner of representing a set of genomic rearrangements corresponds to the type of measurements obtained from DNA sequencing data.

We also define T(*C*), the *terminal* set of *C*, as the set of extremities from *E* appearing in some adjacency in *A*(*C*) but where the corresponding obverse extremity for the interval does not appear in any adjacency in *A*(*C*). Note that an extremity in the terminal set indicates that the associated character (or interval) must appear at the end of the string *C*, and in terms of genomes may be interpreted as the telomere. We define the *extremity permutation π*(*C*) = *π*
_1_
*π*
_2_ … *π_p_
* to be the unique set of extremities appearing in some element of *A*(*C*) after sorting them according to their position in *G*. An extremity permutation *π*(*C*) is *H/T alternating* if its sequence of extremities alternate between being members of the sets *H* and *T*. In such a circumstance we may also refer to *C* as being H/T alternating. Figure 1 shows three examples of strings *C* along with their corresponding adjacency sets *A*(*C*), terminal sets Τ (*C*) and extremity permutations *π*(*C*).

**Figure 1:**
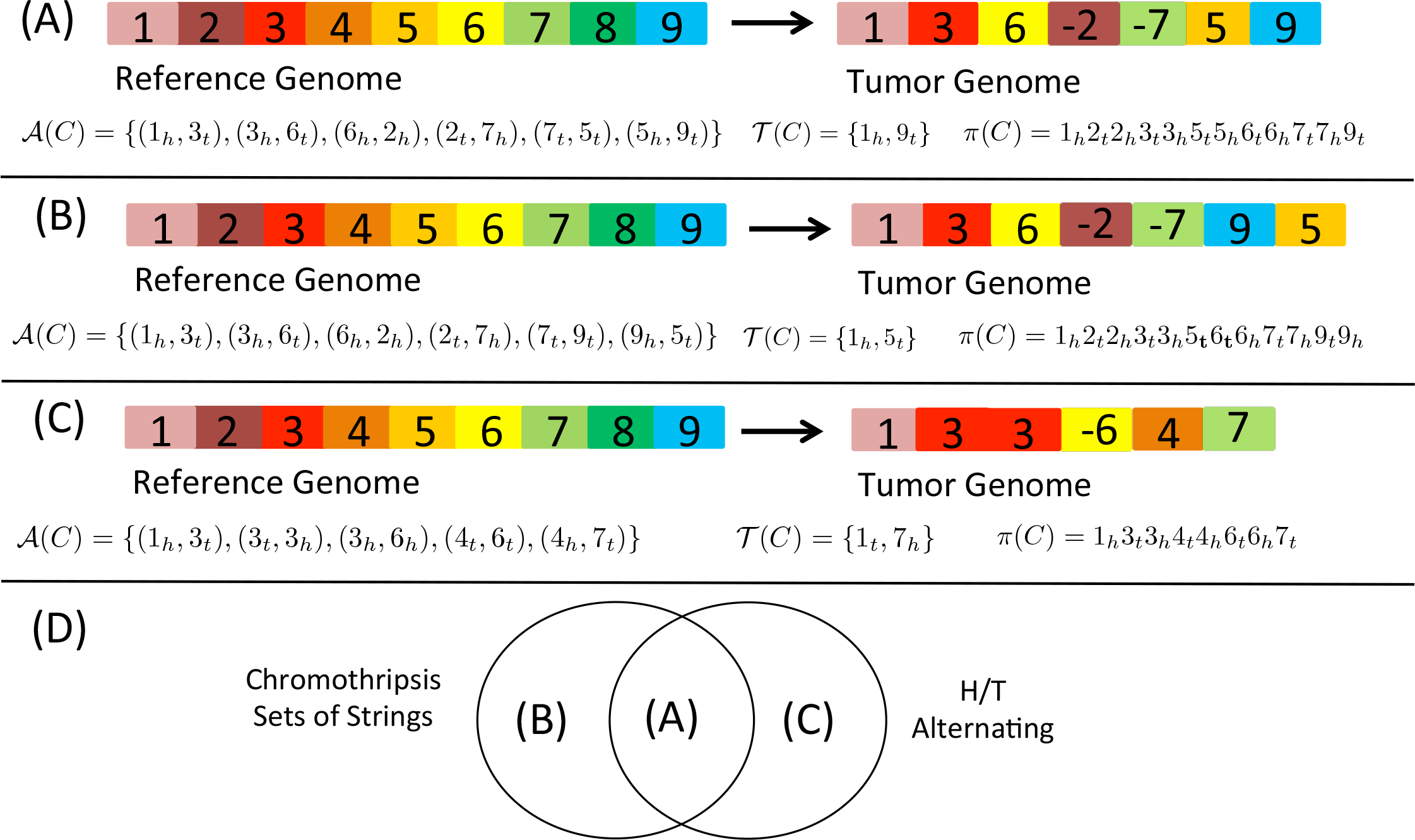
(A)-(C) Three different tumor genomes obtained as a rearrangement of blocks from the reference genome along with their corresponding adjacency sets, terminal sets and extremity permutations. (D) Shows the relationship of the previous three examples in terms of a string *C* being a chromothripsis string or having an extremity permutation *π*(*C*) that is H/T alternating.

### 2.3 H/T Alternating Chromothripsis Strings

We now explore the relationship between our model and the last of the six signatures of chromothripsis suggested by Korbel and Campbell [11] – “the ability to walk the derivative chromosome”. They further describe this signature as defined by an alternating head/tail pattern observed when measured tumor adjacencies are sorted according to their position in the reference genome. In our model, this corresponds to *π*(*C*) being H/T alternating. With our model in place, we can now assess how representative this signature really is of an arbitrary chromothripsis string.

In Theorem 1 we quantify two necessary and sufficient conditions that determine when a chromothripsis string *C* will have an extremity permutation *π*(*C*) that is H/T alternating. In particular, the first condition corresponds to when a chromothripsis event occurs somewhere in the middle of a chromosome, leaving the telomeres in place (Figure 1A), while the second condition corresponds to a degenerate case where both ends of the derivative chromosome originate in a particular configuration from the interior of the chromosome.

#### Theorem 1

*Suppose that C is a chromothripsis string for G. π(C) is H/T alternating if and only if the terminal set j(C) is one of the following:*

1. *T(C) = {π_1_, π_p_} where p = |π(C)|, π_1_
* ∈ *H, and π_p_
* ∈ *T*.
2. *There exists some k such that T(C) = {π_k_, π_k+1_} where π_k_ ∈ T, and π_k+1_ ∈ H*.

*Proof*. Let *C* be a chromothripsis string for *G*.

(*⇒*) Assume that *π*(*C*) is H/T alternating. We will proceed by contradiction. Assume that none of the above conditions about Τ(*C*) are true. In particular, we can also assume that there exists some *k′* ∈ {2, *…, p*} such that *π_k′_ ∈* Τ(*C*) but *π_k′−_
*
_1_, *π_k′_
*
_+1_ ∉ Τ(*C*). Therefore, there must exist some *g, g′* ∈ {1, *…, n*} such that *π_k′−_
*
_1_ = *g_h_
* and *π_k′_
*
_+1_ = *g’_t_
*. However, if *π_k_ H*, this implies that *π*(*C*) does not alternate. Similarly, if *π_k_ Τ*, this implies that *π*(*C*) does not alternate – a contradiction. Hence, one of the above conditions about T(*C*) must be true.

(⇐) We will consider each possible telomere set Τ(*C*) separately and show that for each it is true that *π*(*A*(*C*)) is alternating.

Assume that Τ(*C*) = {*π*
_1_, *π_p_
*} where *p* = |*π*(*C*)|, *π*
_1_ ∈ *H*, and *π_p_ T*. We will proceed by contradiction. Assume that *π*(*C*) is not alternating. This implies (without loss of generality) that there exists some *k ∈* {1, *…, p −* 1} and *g, g^’^ ∈* {1, *…, n*} such that *π_k_
* = *g_h_, π_k_
*
_+1_ = *g^’^
_h_
* (that is *π_k_, π_k_
*
_+1_ *∈ H*). This implies that *g’_t_ ∉ π*(*C*) and therefore *g’_h_
* = *π_k_
*
_+1_ *∈* Τ(*C*). And since *k* + 1 *>* 1, it must be the case that *g′_h_
* = *π_k_
*
_+1_ = *π_p_
*, therefore contradicting our assumption that *π_p_ ∈ T*. The argument for *π_k_
* = *g_t_, π_k_
*
_+1_ = *g’_t_
* is similar. Hence, *π*(*C*) must be H/T alternating.

Assume there exists some *k* such that Τ(*C*) = *π_k_, π_k_
*
_+1_ where *π_k_
* ∈ *T*, and *π_k_
*
_+1_ ∈ *H*. We will proceed by contradiction. Assume that *π*(*C*) is not alternating. This implies (without loss of generality) that there exists some *k’ ∈* {1, *…, p −* 1} and *g, g’ ∈* {1, *…, n*} such that *π_k′_
* = *g_h_, π_k′_
*
_+1_ = *g’_h_
* (that is *π_k′_, π_k′_
*
_+1_ *∈ H*). This implies that *g’_t_ ∉ π*(*C*) and therefore *g’_h_ ∈* Τ(*C*). There are only two possible values of *k^’^
* such that *π_k_′*
_+1_ *∈* Τ(*C*). The first possibility is that *k’* = *k −* 1. If *k’* = *k −* 1, then *π_k_
* = *g’_h_
*, a contradiction with our assumption that *π_k_ ∈ T*. The second possibility is that *k^’^
* = *k*. If *k’* = *k*, then *π_k_
* = *g_h_
*, a contradiction to our assumption that *π_k_ ∈ T*. The argument for *π_k′_
* = *g_t_, π_k′_
*
_+1_ = *g’_t_
* is similar. Hence, *π*(*C*) must be H/T alternating.

It is important to note that the cases detailed in Theorem 1 do not include the case where a chromothripsis event includes one telomere and the final segment joined to form the end of the derivative chromosome is different than the original telomere (Figure 1B). This is potentially an observation of significance as recent studies have suggested that chromothripsis may arise as the result of telomere crisis [13]. Thus, *even in the absence of noise*, a chromothripsis string may not be H/T alternating. Furthermore, a linear string *C* that is *not* a chromothripsis string may still have an extremity permutation *π*(*C*) that is H/T alternating (Figure 1C).

### 2.4 Fraction of Chromothripsis Strings that are H/T alternating

In the previous section we demonstrate that not all chromothripsis strings *C* have extremity permutations *π*(*C*) that are H/T alternating and categorize the subset that does exhibit this property. If the vast majority of chromothripsis strings exhibit this signature, then it may still be useful for categorizing chromothripsis. We therefore derive a formula for the exact fraction of chromothripsis strings that are H/T alternating in Theorem 2 (see Appendix for proof).

#### Theorem 2

*The fraction of chromothripsis strings of length m derived from a reference genome G composed of n intervals with π(C) that is H/T alternating is* 
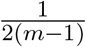
.

Theorem 2 tells us that the fraction of chromothripsis strings that are H/T alternating depends only on the number m of characters in the chromothripsis string and not the original number of intervals n. More importantly, as m increases this fraction decreases quickly (Figure 2). For example, only 5.6% of chromothripsis strings containing 10 intervals are H/T alternating. Thus, in its current stringent form, the H/T alternating property does not seem well suited as a signature of chromothripsis.

**Figure 2:**
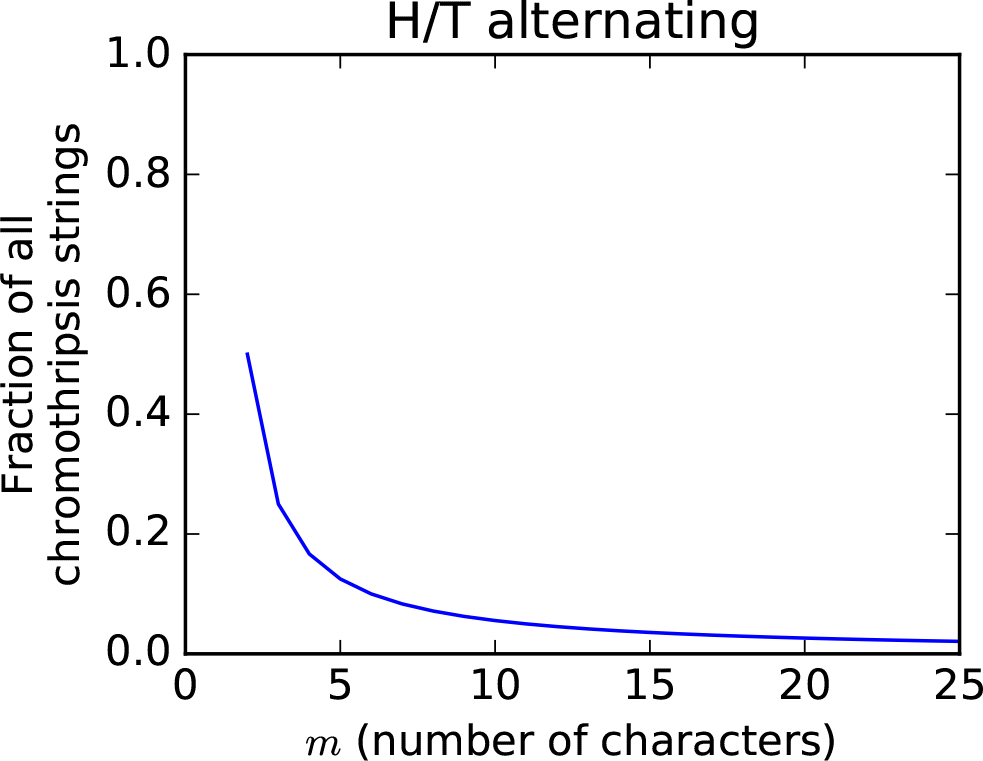
The fraction of chromothripsis strings *C* that are H/T alternating according to Theorem 2.

### 2.5 An Alternative Signature: H/T Alternating Fraction

Treating H/T alternating as a binary feature that a chromothripsis string *C* either exhibits or it doesn’t, prevents us from capturing any information about how close or far *C* is from H/T alternating. For example, consider the following two sequences of H/T membership that an extremity permutation of length 8 might exhibit: HTHTHTHH and HHHHTTTT. Neither one is H/T alternating, but the former is much closer to H/T alternating than the later. Thus, it would be useful to have a measure that allows us to capture this type of more nuanced information.

We therefore define the *H/T alternating fraction* of a linear string *C* or *AF* (*C*) as the fraction of adjacent pairs of characters in *π*(*C*) = *π*
_1_
*π*
_2_ … *π_p_
* that have one character from the set of head extremities *H* and the other from the set of tail extremities *T*. Formally, *alt*(*π_i_, π_j_
*) tells us whether or not a selected pair of characters *π_i_
* and *π_j_
* include one character from *H* and one from *T*.

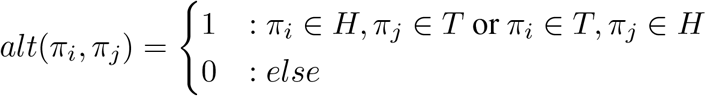

Given a linear string *π*(*C*) = *π*
_1_
*π*
_2_ … *π_p_
* of length *p* we calculate *AF* (*C*) according to the following equation:

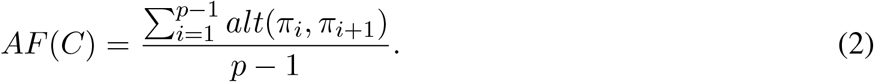

The H/T alternating fraction is a measure that allows us to precisely capture the more nuanced information that was lacking from the more stringent definition of H/T alternating. A chromothripsis string *C* that is H/T alternating will have *AF* (*C*) = 1.0 and chromothripsis string *C* that lacks the H/T alternating property (due to the shattering occurring at the telomere) will still have a robust H/T alternating fraction because at most two adjacent pairs of extremities (representing the two telomeres) will disrupt the alternating pattern.

### 2.6 Distinguishing Between Simultaneous and One-off Events

In the previous sections we showed that the H/T alternating fraction allows for a more robust version of the signature originally suggested by Korbel and Campbell [11]. Here we consider whether this signature is able to distinguish between one-off events such as chromothripsis and step-wise events.

First we note that a cancer genome *C* derived from simple, non-overlapping rearrangements may in fact result in a rearrangement string that is a chromothripsis string. For example, consider a reference genome *G* = (1, 2, 3) and a cancer genome *C* = (1,*–*2, 3) having undergone an inversion of interval 2. In this instance *AF* (*C*) = 1.0. In the most extreme case, a non-rearranged cancer genome *C* = (1, 2, *…, n*) would also be a chromothripsis string and have *AF* (*C*) = 1.0. So while such genomic rearrangements are technically possible from a chromothripsis event, these configurations seem unlikely to have arisen that way, and we would likely not want to classify them as the result of chromothripsis. One key distinction in these cases is that these simple rearrangements do not include any of the other traditional signatures of chromothripsis, including overlapping rearrangements. Therefore, we will focus on the utility of the H/T alternating fraction when considering overlapping rearrangements where the distinction of chromothripsis versus a step-wise sequence of events is much less obvious.

Next, we consider a different situation. Consider a reference genome *G* = (1, 2, 3, 4, 5) and a cancer genome *C* = (1, 2, 3, 2, 5) obtained through a tandem duplication of intervals 2, 3 followed by an overlapping deletion of intervals 3, 4. In this instance, *C* is not a chromothripsis string as it contains a duplication. If we observed the exact sequence of intervals in *C* we would have *A*(*C*) = {(1_h_, 2_
*t*
_), (2_h_, 3_
*t*
_), (3_h_, 2_
*t*
_), (2_h_, 5_
*t*
_)}, *π*(*C*) = 1_
*h*
_2_
*t*
_2_
*h*
_3_
*t*
_3_
*h*
_5_
*t*
_ and *AF* (*C*) = 1.0. This would seem problematic for our goal of using H/T alternating fraction to distinguish between one-off and simultaneous events. However, we make the important observation that we don’t actually observe the sequence of intervals in a cancer genome. Instead we observe the set of novel adjacencies (not occurring in the reference genome) between regions of the genome and it is these adjacencies that define the set of genomic intervals used in the derivative chromosome. So, for the above example we would only observe *A*(*C*) = {(3_h_, 2_
*t*
_), (2_h_, 5_
*t*
_)} which leads to *π*(*C*) = 2_
*t*
_2_
*h*
_3_
*h*
_5_
*t*
_ which has lower *AF* (*C*) = 0.66. We expect similar drops in *AF* (*C*) values in other situations with overlapping rearrangements as well. Thus, the H/T alternating fraction would appear to be the most useful when considering sets of overlapping aberrations, and while certainly not a perfect classifier between one-off and simultaneous events, it has the potential to be useful in some situations.

### 2.7 Extending to Multiple Chromosomes

Thus far we have presented this work in terms of a unichromosomal genome *G*. Extending to the case of a multi-chromosomal genome is straightforward. This is done by considering the set of extremities originating from each chromosome separately. For example, if the extremity permutation for each chromosome is H/T alternating, then entire set is H/T alternating. Similarly, the H/T alternating fraction is computed by considering only adjacent pairs of extremities appearing within the same chromosome.

## 3 Results

We demonstrate the usefulness of the H/T alternating fraction measure on both simulated and real DNA sequencing data.

### 3.1 Simulations

We simulate data by shattering a reference genome into *n* blocks and then create random chromothripsis genomes that have *m* novel adjacencies between those blocks. We introduce noise into the set of adjacencies *A*(*C*) by either removing true adjacencies or adding extraneous adjacencies to the set (see Appendix for further details).

For different values of *m* we verified that our observed fraction of chromothripsis genomes that are strictly H/T alternating matched well with our theoretically derived fraction (Theorem 2). These fractions are quite low to begin with (for example only 3.5% of 10,000 random genomes with *m* = 15 adjacencies are H/T alternating) and this fraction becomes even smaller as noise is added (see Appendix Figure C.1). Therefore, we restricted our attention to chromothripsis genomes that are originally H/T alternating, and even in that instance we observe that the H/T alternating signature degrades quickly with noise (see Appendix Figure C.2). For example, less than 3% of 10,000 simulations with *m* = 15 novel adjacencies are H/T alternating when just a single adjacency is either added or removed. We do note that Korbel and Campbell [11] indicate that the version of this signature they propose is dependent on observing all adjacencies. In contrast, we show that the H/T alternating fraction measure *AF* (*C*) is much more robust to noise (Figure 3). For example, over simulations with *m* = 35 novel adjacencies in the chromothripsis genome, adding or removing up to 4 adjacencies produces an alternating fraction *AF* (*C*) above 0.9 and 0.87, respectively, in all 10,000 random trials.

**Figure 3:**
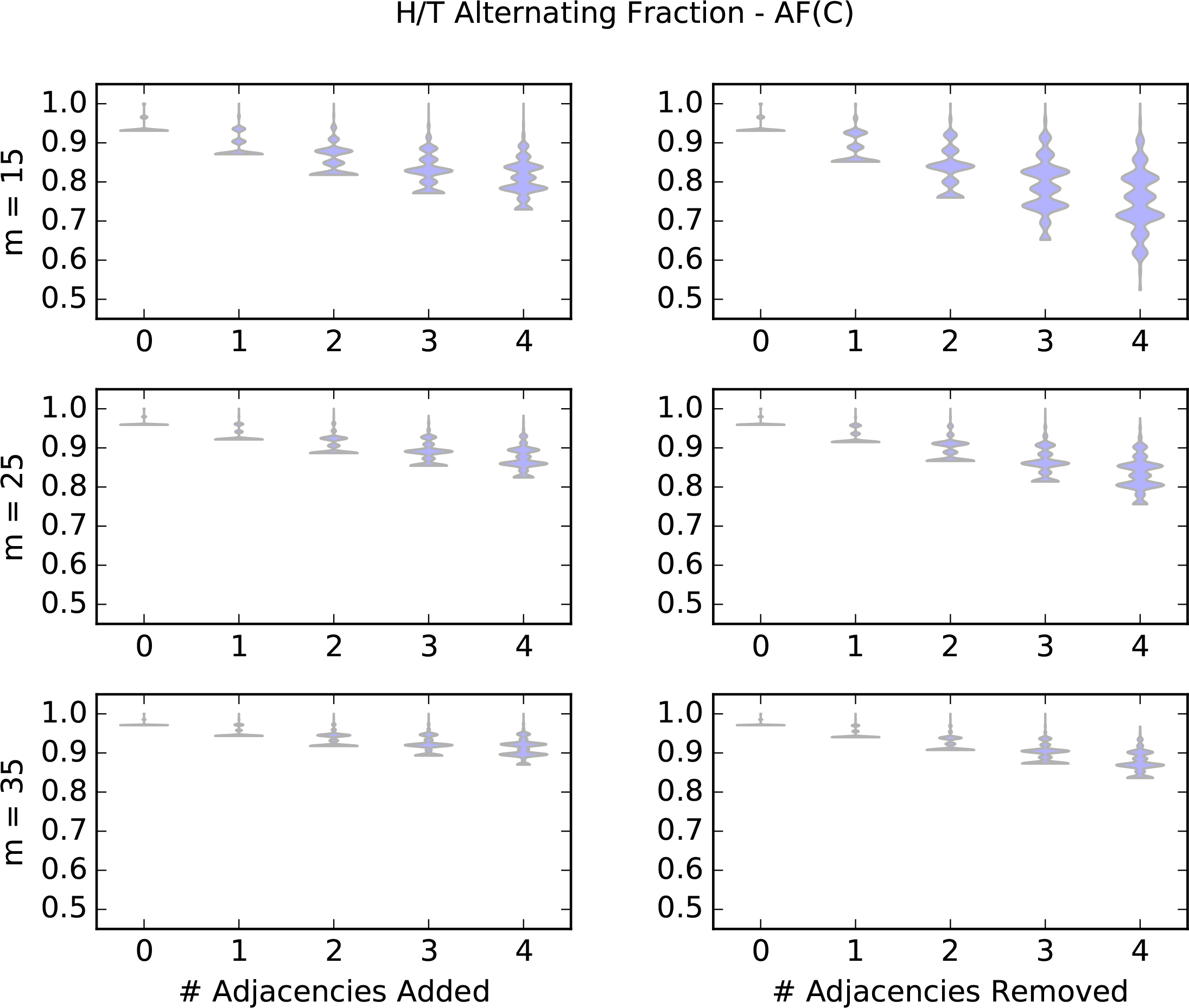
Violin plots over 10,000 randomly generate chromothripsis genomes (with *m* novel adjacencies) showing that the alternating fraction *AF* (*C*) measure remains high even when noise is incorporated by randomly adding and removing adjacencies.

The H/T alternating fraction measure is more useful if it can distinguish chromothripsis genomes from genomes that have undergone a sequential accumulation of events. Therefore we simulated genomes that have undergone a sequential accumulation of random events including deletions, tandem duplications and inversions. We created these genomes by randomly adding such events to a reference genome until the number of novel adjacencies was equal to or exceeded a specified value *m* (see Appendix for further details). We compare the observed distribution of H/T alternating fractions for chromothripsis genomes and sequential genomes (Figure 4) and find that in the case of noise-free data, the H/T alternation fraction or *AF* (*C*) value for chromothripsis genomes is much higher than for the sequential genomes. We also created simulated data where where mutations occur as part of an evolutionary branching process instead of a single step-wise accumulation of mutations (see Appendix for further details) and found the results to be very similar to those presented here (Appendix Figure C.3). It is important to note that the observed difference in H/T alternating fraction between the one-off genomes and the step-wise genomes shown in Figure 4 will diminish as noise is added into the data. In particular, the corresponding panel in Figure 3 with *m* = 25 shows that noise will lower the H/T alternating fraction for the one-off genomes, but will still remain largely above what we observe here for the step-wise genomes. Thus, these results indicate that the H/T alternating fraction does indeed capture some quantifiable aspect about the presence of chromothripsis. While certainly not perfect – some sequentially generated genomes may also have a high H/T alternating fraction – the signature appears applicable in some instances.

**Figure 4:**
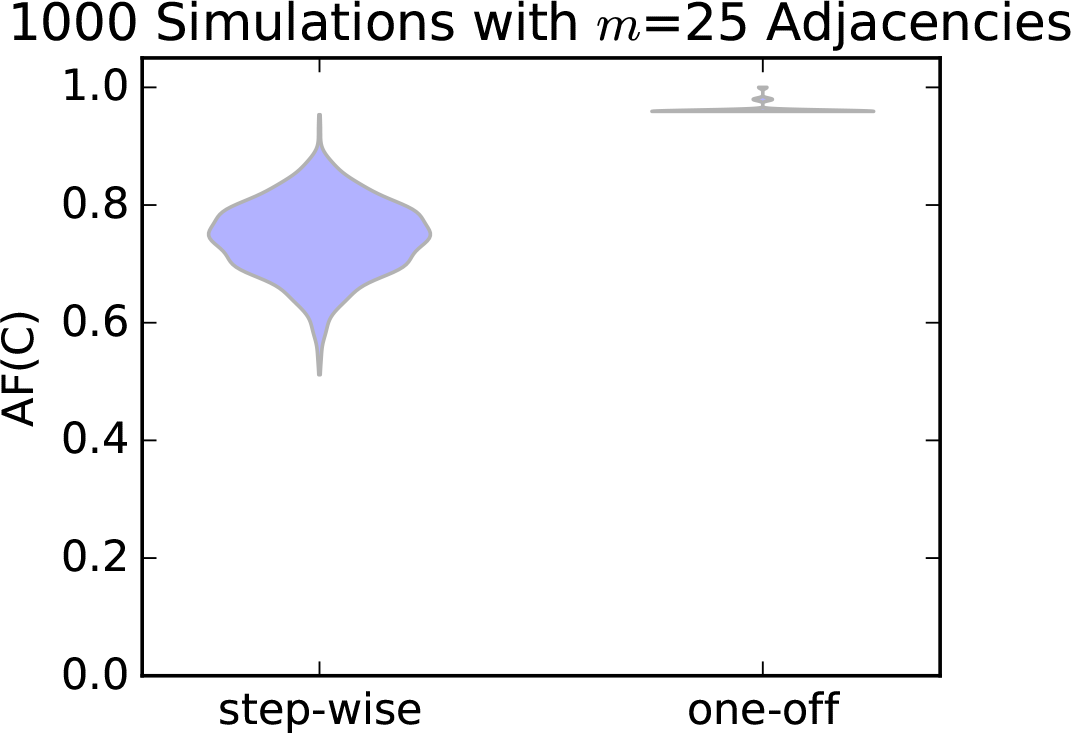
In error-free simulations, we observe that the H/T alternating fraction *AF* (*C*) measure is much higher for one-off (chromothripsis) genomes than genomes that have undergone a step-wise sequence of events and have the same number of novel adjacencies.

### 3.2 Real Data

We apply our H/T alternating fraction measure to a dataset of 64 genomes representing seven tumor types from the The Cancer Genome Atlas (TCGA) that were previously analyzed for chromothripsis by Malhotra et al. [14]. In particular, they identify 154 sets of observed adjacencies that they classify as either one-off (chromothripsis) events (97 sets of adjacencies) or as step-wise (57 sets of adjacencies). We compute the H/T alternating fraction across each of these sets (Figure 5A) and find that the mean of the one-off set is 0.689 and the mean of the step-wise set is 0.602 with a difference in means of 0.087. We use a permutation test with 100,000 permutations to assess the statistical significance of the observed difference in means between the two sets, resulting in a permutation distribution with mean of 7.5 × 10^
*−*5^ and standard error of 0.032, and find that the H/T alternating frequency of the one-off events is statistically higher than the step-wise events (*p* = 0.00265). We also compute a bootstrap-t confidence interval over 10,000 replicates and determine that with 95% the average one-off H/T alternating fraction is between 0.029 and 0.15 units larger than the average step-wise H/T alternating fraction. For additional statistical analysis of the differences between these distributions, see the Appendix.

**Figure 5:**
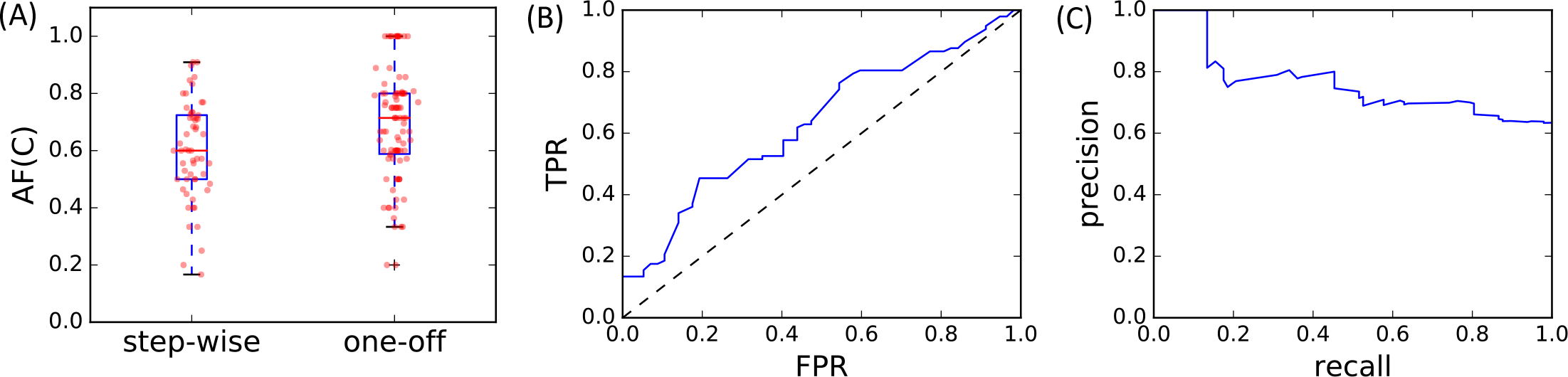
(A) Sets of adjacencies previously classified by Malhotra et al. [14] as one-off (chromothripsis) have a statistically higher observed H/T alternating fraction *AF* (*C*) that those sets classified as step-wise (*p* = 0.00265 with a permutation test). (B) ROC curve (*AU C* = 0.63) and (C) precision-recall curve when events labeled as one-off are considered the positive class and the threshold for classifying as one-off orstep-wise based on *AF* (*C*) is varied.

We also analyze the use of different thresholds *t* of the H/T alternating fraction, or *AF* (*C*), to classify a sample as one-off (*AF* (*C*) ≥ *t*) or step-wise (*AF* (*C*) < *t*). In particular we see that *AF* (*C*) serves as a modest classifier by considering the associated ROC curve (*AU C* = 0.63) and precision-recall curve (*AU C* = 0.76) as shown in Figures 5B and 5C respectively. While these results are not strong enough to support the use of the H/T alternating fraction as a classifier in isolation, they do indicate that this signature does provide some signal and may therefore lead to improved results when used in conjunction with other signatures to do classification.

We note that our computations of H/T alternating fraction also allows us to determine which, if any of these samples, exhibit the strict H/T alternating criterion as they would have a value of *AF* (*C*) = 1.0. We find that only 13 of the 154 sets of adjacencies have *AF* (*C*) = 1.0, all of which were originally classified as one-off. However, we also note that these sets contain relatively few breakpoints, only 6 or 8 in each one, the smallest values contained in any sets in the entire dataset.

Weinreb *et al*. [22] later re-analyzed this dataset and identified some sets of adjacencies that may have originally been misclassified. This includes lung squamous cell carcinoma sample LUSC-11 (adjacency set chain-5) which Malhotra *et al*. originally classified as step-wise but Weinreb *et al*. suggest that it may indeed be the result of a one-off event like chromothripsis. We find a relatively high H/T alternating fraction value of 0.73 for this sample. While this is not the highest H/T alternating fraction found for any set of rearrangements originally labeled as step-wise (in fact it is the 12th highest), this set of rearrangements contains 84 breakpoints along a single chromosome, which is many more than the 11 step-wise rearrangement sets with higher H/T alternating fractions that have on average only 14.7 breakpoints. Observing a high H/T alternating fraction with more breakpoints may provide additional confidence that the pattern is not due to chance. We interpret these results as further supporting evidence that this sample may indeed have originally been misclassified.

## 4 Discussion and Conclusion

Determination of a rigorous signature that may be useful for identification of a one time event, such as chromothripsis, has proven to be a challenging task [7]. A number of different criteria for inference of chromothripsis have been proposed [21] including a list of six signatures proposed by Korbel and Campbell [11]. While some of these signatures (e.g clustering of breakpoints, oscillation of copy-number states) have been widely used in practice [14, 6, 19] others have been less studied. In this work we provide a rigorous formulation and analysis of the “ability to walk the derivative chromosome” signature proposed by Korbel and Campbell [11], which we refer to as H/T alternating. In particular, we have shown that this signature, as originally envisioned, may not always be present in a chromothripsis genome and we provide a precise quantification of under what circumstances it would be present. We then propose a variation on this signature, the *H/T alternating fraction*, which allows us to measure to what degree the H/T alternating property, originally defined by [11], is present throughout the genome. We apply this measure to a previously analyzed dataset and find that sets of rearrangements previously classified as one-off (chromothripsis) have a statistically higher H/T alternating fraction than those classified as step-wise. Thus, indicating that the H/T alternating fraction may be an indicative measure of rearrangements obtained simultaneously as opposed to sequentially.

While many studies have investigated the occurrence of chromothripsis in different tumor types, the relative prevalence of the phenomenon remains unknown [20]. Some studies have attempted to estimate the rate of chromothripsis by considering large datasets utilizing existing methods or signatures to identify likely chromothripsis candidates. For example, Cai *et al*. [2] identified 918 cancer samples with chromothripsis-like patterns from a dataset of *>* 22,000 cases. This yields a relatively low prevalence rate of only 4.2%. This is similar to the 2-3% prevalence rate estimated in the original Chromothripsis publication [21]. However, despite low prevalence across all tumor types, the ability to accurately determine these instances has important potential impact as samples labeled as chromothripsis are often associated with poor patient outcome [10, 16]. Furthermore, the estimated rate of chromothripsis is much higher in some types of cancer. For example, Stephens *et al*. [21] estimated a rate of 25% in bone cancers. Thus, improved methods for identifying genomes that may have undergone a catastrophic event like chromothripsis is an important and necessary step in the search to better understand and treat cancer patients.

There are a number of avenues for further investigation relating to this work. In particular, we have demonstrated the applicability of the H/T alternating fraction on one curated dataset where sets of rearrangements had already been classified as either one-off (chromothripsis) or step-wise. Many other chromothripsis studies do not include sets of rearrangements classified as step-wise in addition to those classified as chromothripsis, making it difficult to assess how well this signature generalizes to other real datasets. We did calculate the H/T alternating fraction for 15 additional genomes classified as chromothripsis by two additional studies by Stephens *et al*. [21] and Rausch *et al*. [19]. But since both of these studies did not include genomes having undergone similar processing and called as step-wise, it is difficult to appropriately assess the meaning of their H/T alternating fractions. To this end, we did add these genomes to the set classified by [14] as one-off and compared them to the set classified as step-wise, and still find a significant difference in the H/T alternating fraction of the two groupings (*p* = 0.0036 with a permutation test comparing the difference of the means of the two groupings). Further study of this signature applied to new datasets as they become available may yield further insight into its usefulness.

Another important area for further investigation is how this signature may be the most useful in practice for classifying rearrangements as sequential or simultaneous. In this work we have focused mainly on analyzing the “ability to walk the derivative chromosome signature” proposed by [11] and one variation of this signature in a largely proof-of-concept context. Given that we have shown that the H/T alternating fraction degrades with noise and is dependent on the number of adjacencies involved in the rearrangement, we suspect that this signature may be most useful when used in combination with other proposed signatures of chromothripsis, such as clustering of breakpoints or oscillation of copy number, rather than in isolation. Exploration of how to integrate this signature with other signatures in order to detect chromothripsis events, perhaps in the context of new or existing tools such as ShatterProof [6], is left as future work.

## A Full Proofs Omitted in the Main Text

(**Main Text) Theorem 2.** *The fraction of chomothripsis strings of length m derived from a reference genome G composed of n intervals with π(C) that is H/T alternating is* 
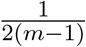

In order to prove this theorem we will first need to build up some more notation and observations.

First, we calculate *|G(m, n)|*, the total number of chromothripsis strings of *m* blocks given a reference genome of n blocks. This is straightforward using the following equation.

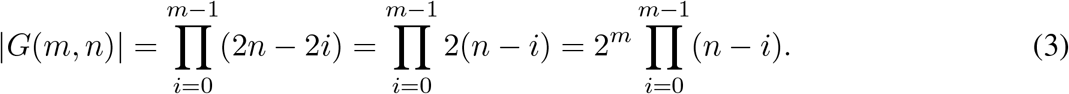

Next we calculate *|A(m, n)|*, the number of chromothripsis strings of *m* blocks given a reference genome of *n* blocks that are H/T alternating. This computation utilizes the fact that the two cases from Theorem 1 are mutually exclusive and thus the number of instances for each one can be counted separately and then summed together. Furthermore, the selection of any two blocks (telomeres) in a reference genome *G = 1 … n* defines a partition of the remaining *n* – 2 blocks into two sets: (1) blocks that lie between the two chosen telomere blocks in *G;* and (2) blocks that lie outside the chosen telomere blocks in *G*. In Case 1 from Theorem 1 all non-telomere blocks in the derivative chromosome lie in between the telomeres and in Case 2 they must lie outside the telomeres. For each possible number of blocks that fall between or outside the telomeres (ranging from *m*−2 up to *n*−2) in *G* we can explicitly count the number of potential telomere pairs and configurations the *m* − 2 other blocks. These observations (along with some algebra) allow us to derive the following formula for *|A(m, n)|*.

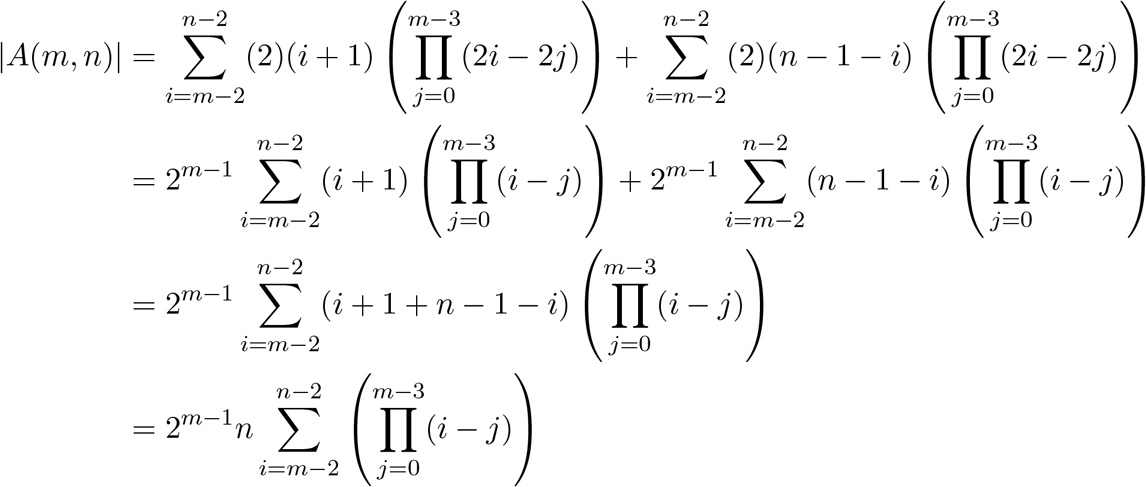

Lastly, we prove Lemma A.1 which is used in the proof for (Main Text) Theorem 2.

**Lemma A.1**

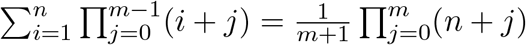

*Proof*. We use proof by induction on the variable *n*. We start with the base case *n* = 1.

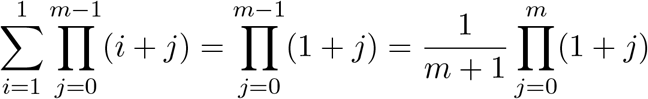

We now assume the property holds for values up to *n* − 1 and want to prove for generic *n*.

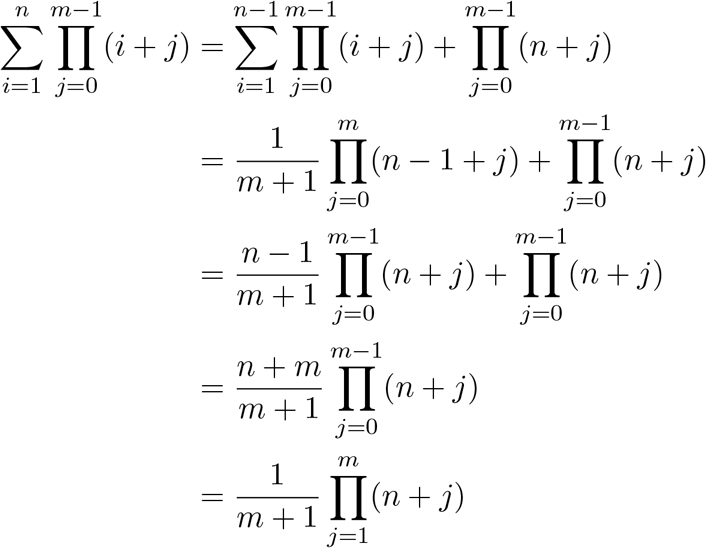

Therefore we have proven the Lemma for general *n*.

Now we can provide a full proof of (Main Text) Theorem 2

*Proof*. The fraction of chromothripsis strings of length *m* derived from a reference genome *G* composed of *n* intervals that are H/T alternating is just the the number of such chromothripsis strings that are H/T alternating divided by the total number of such chromothripsis strings.

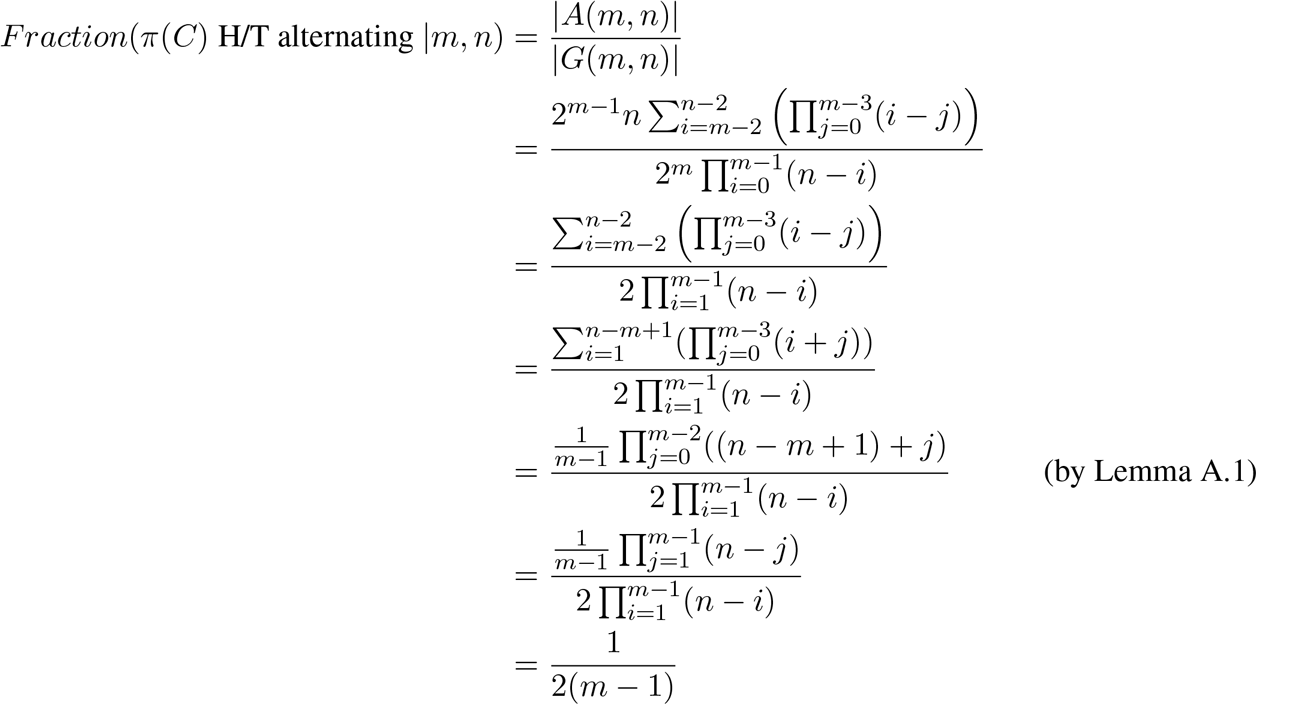

## B Simulation Details

In this section we provide further details on how we simulate data.

### B.1 Chromothripsis Genomes

When simulating genomes that have undergone a chromothripsis event, we aim to create a genome that has a specified number *m* of novel adjacencies. We do this with the following procedure. We first construct a reference genome that is partitioned into a specified number of blocks *n*. We then create a random signed permutation of these blocks. We then construct the chromothripsis genome, as a sequence of these blocks by starting at the beginning of the permutation and counting the number of novel adjacencies we observe in the sequence as we move forward through the permutation. One we have a sequence of blocks with *m* novel adjacencies (or we have run out of blocks), we return that set of blocks as the chromothripsis string *C*. More specifically, since the *AF* (*C*) value is computed using observed adjacencies, we return the set of *m* novel adjacencies *A*(*C*) observed when traversing the permutation.

We add noise into this data by adding and removing random adjacencies from *A*(*C*). Removal is done by randomly selecting an adjacencies in *A*(*C*) and removing it from the set. In order to add random adjacencies, we first create a set of possible adjacencies to add. We create this set by looking at the permutation of all *n* blocks used to create the chromothripsis genome *C* and add to that set every novel adjacency of blocks that was not included in the random chromothripsis genome. We then select adjacencies from this set to add to *A*(*C*).

### B.2 Step-wise Mutations

When simulating genomes that have undergone a sequential accumulation of events we begin by breaking the reference genome into a specified number of blocks *n*. We then add random rearrangement events that are either deletions, duplications or inversions to the genome. The position of a rearrangement event is randomly determined and its size is also randomly determined. We do however restrict the maximum size of a rearrangement event. We want to restrict the size of events in order to obtain a genome that more realistically resembles what might be seen in real data. For example, if we allowed deletion events to contain any number of blocks, often times we may end up deleting large portions of a genome (or nearly all of it) leading to simulations were we are unable to obtain the desired number of adjacencies. For the simulations presented in this work we set *n* = 100 and restrict the size of events to be no larger than 5 genome blocks/segments.

### B.3 Step-wise Mutations along with Branching Evolutions

We also create simulations that incorporate evolutionary branching processes, where each branch may contain a different collection of sequentially obtained mutations. We begin by breaking the reference genome into a specified number of blocks *n* and start with a single, non-rearranged genome in our set of genomic populations. We then pick a single genome in our set of existing populations that we will modify by adding to it a single randomly generated aberration (using the approach described above). However, before adding this rearrangement to the selection population we first decide if this rearrangement will represent a branch in the evolutionary history of this genome. We randomly add a branch with probability *α* and this consists duplicating the selected genomic population in the set of existing populations and then adding a random rearrangement to one of the copies of this population. We then consider the entire set of novel rearrangements existing with this population of genomes when constructing *π*(*C*) and determining its H/T alternating fraction. For the simulations presented in this work we set *n* = 100, restricted the size of events to be no larger than 5 genome blocks/segments and used *α* = 0.2. Once *m* = 25 novel aberrations were created within the set of populations we stopped the simulation procedure. With this value of *α* and *m* the expected number of tumor populations per simulation is 6 (the original population plus 5 branches).

## C Additional Results

### C.1 Statistical Analysis of Malhotra *et al*. Data

We performed additional statistical analysis of the 154 sets of adjacencies reported by Malhotra *et al*. [14] as either one-off (chromothripsis) events (97 sets of adjacencies) or as step-wise (57 sets of adjacencies). We compute the H/T alternating fraction across each of these sets use a Mann-Whitney test to determine that the alternating frequency of the one-off events is statistically higher than that step-wise events (*p* = 0.0028), but with a small to medium effect size (*r* = 0.22).

**Figure C.1:**
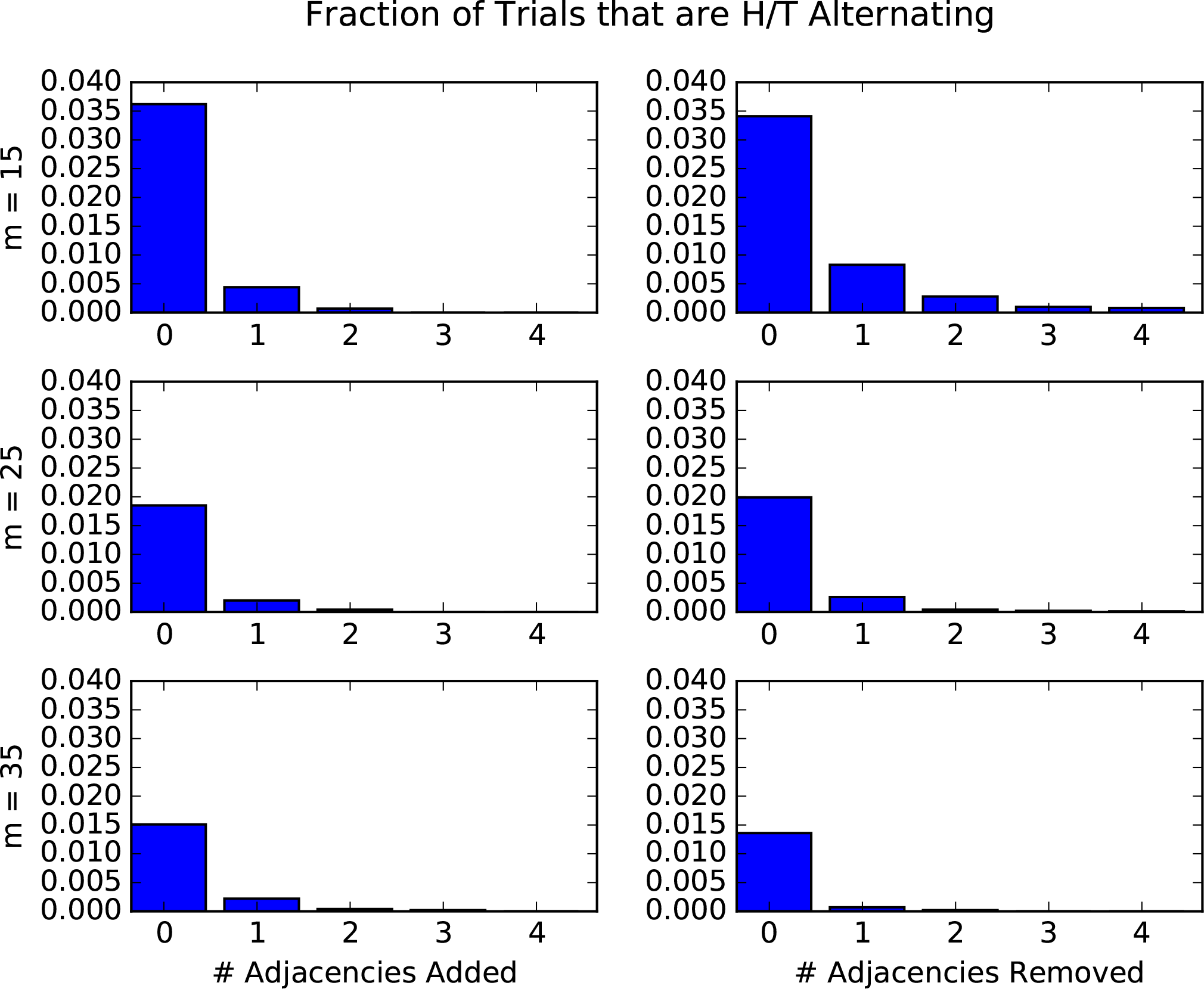
For each value of *m* (number of novel adjacencies in the chromothripsis genome) we created 10,000 random chromothripsis genomes. We then incorporated noise into the data by randomly adding and removing adjacencies and then determined what fraction of the resulting datasets were H/T alternating.

**Figure C.2:**
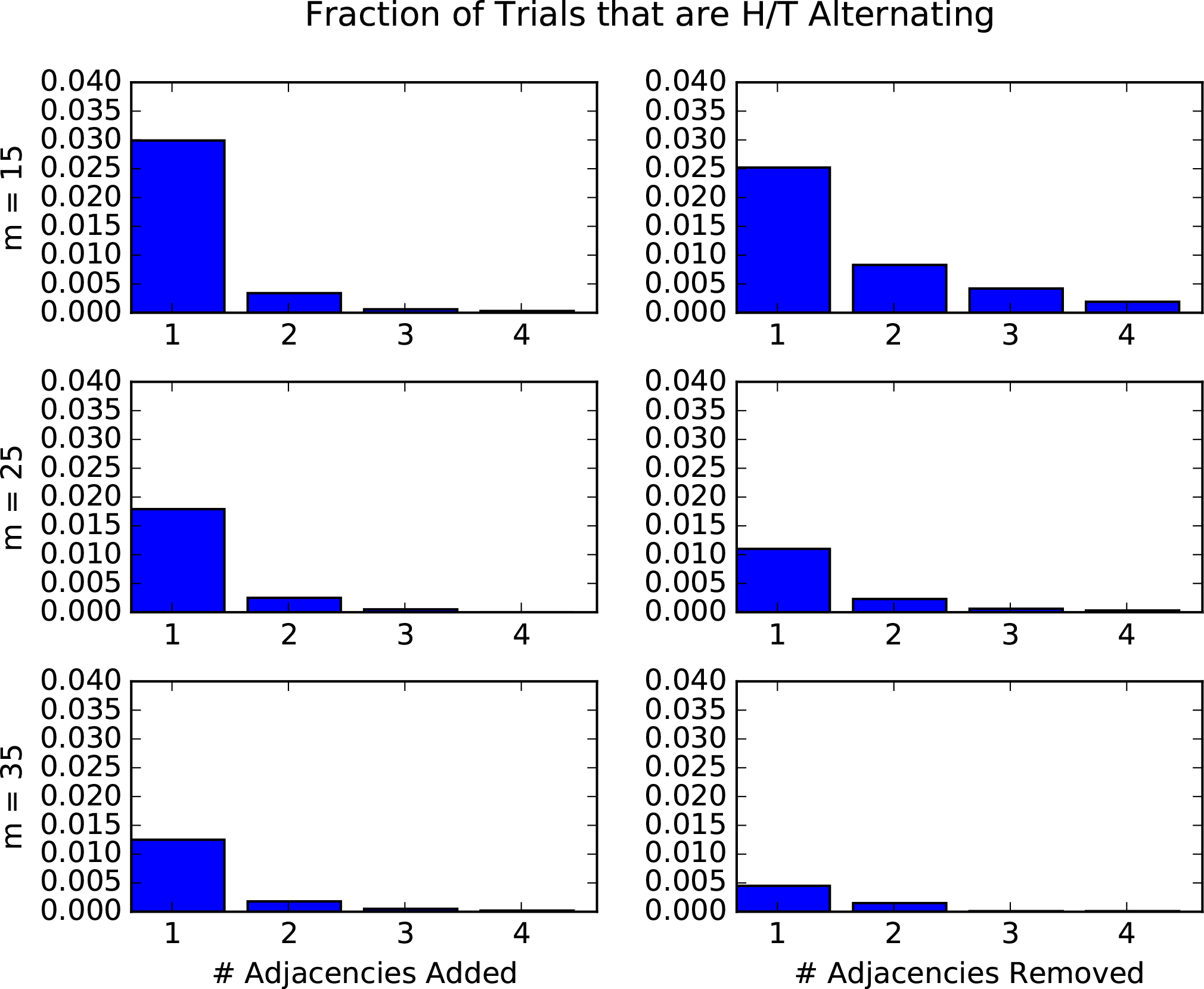
For each value of *m* (number of novel adjacencies in the chromothripsis genome) we created 10,000 random chromothripsis genomes that were initially H/T alternating. We then incorporated noise into the data by randomly adding and removing adjacencies and then determined what fraction of the resulting datasets were H/T alternating.

**Figure C.3:**
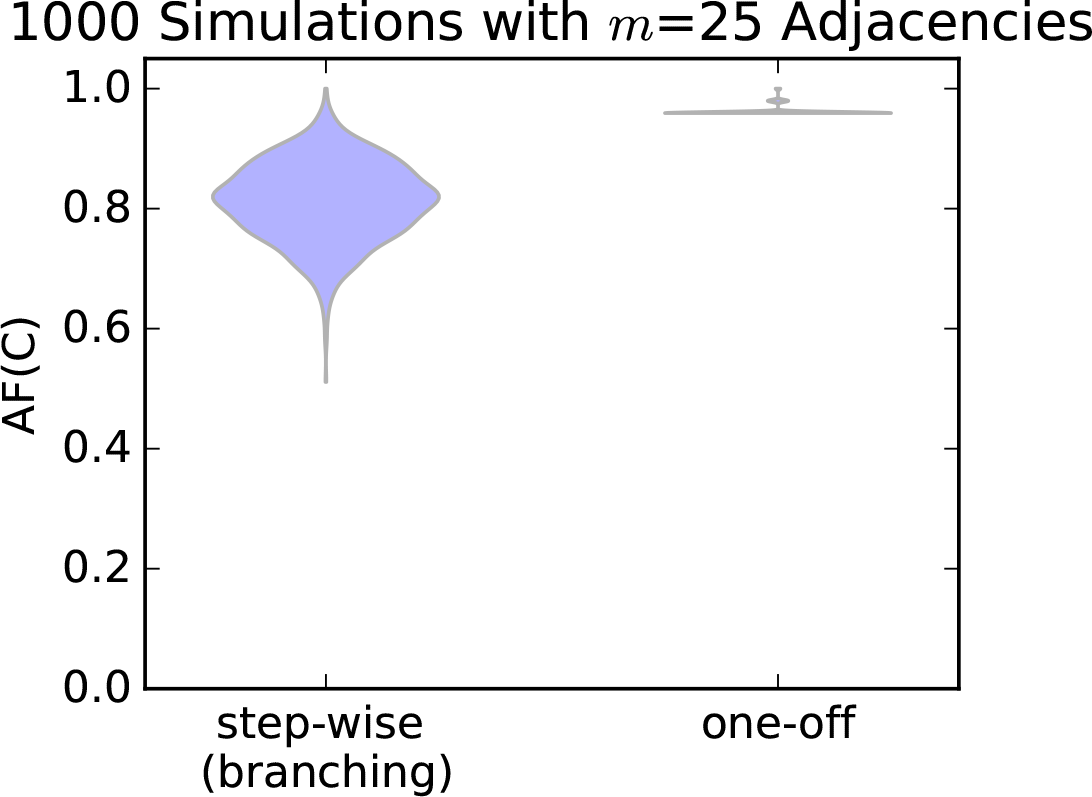
In error-free simulations, we observe that the H/T alternating fraction *AF* (*C*) measure is much higher for one-off (chromothripsis) genomes than genomes that have undergone step-wise sequences of events occurring during branching evolution and have the same number of total novel adjacencies.

